# Senescent cells in Giant Cell Arteritis have inflammatory phenotype participating in tissue injury via IL-6 dependent pathways

**DOI:** 10.1101/2023.05.19.541093

**Authors:** D Veroutis, OD Argyropoulou, AV Goules, K Kambas, DA Palamidas, K Evangelou, S Havaki, A Polyzou, E Xingi, E Karatza, K Boki, A Cavazza, C Kittas, D Thanos, C Ricordi, C Marvisi, F Muratore, E Galli, S Croci, C Salvarani, VG Gorgoulis, AG Tzioufas

**Affiliations:** Molecular Carcinogenesis Group, Department of Histology and Embryology, Medical School, National and Kapodistrian University of Athens, Athens, Greece; Department of Pathophysiology, School of Medicine, National and Kapodistrian University of Athens, Greece; Research Institute for Systemic Autoimmune Diseases, Athens, Greece; Laboratory of Molecular Genetics, Department of Immunology, Hellenic Pasteur Institute, Athens, Greece; Light Microscopy Unit, Hellenic Pasteur Institute; Second Propaedeutic Department of Surgery, National and Kapodistrian University of Athens, Laikon General Hospital; Rheumatology Unit, Sismanoglion Hospital, Athens, Greece; Pathology Unit, Azienda Unità Sanitaria Locale-IRCCS Di Reggio Emilia, Reggio Emilia, Italy; Faculty of Health and Medical Sciences, University of Surrey, Guildford, UK; Center of Basic Research, Biomedical Research Foundation of the Academy of Athens, Athens, Greece; Unit of Rheumatology, Azienda Unità Sanitaria Locale-IRCCS di Reggio Emilia, Reggio Emilia, and University of Modena and Reggio Emilia, Reggio Emilia, Italy; Unit of Clinical Immunology, Allergy and Advanced Biotechnologies, Azienda Unità Sanitaria Locale-IRCCS di Reggio Emilia, Italy; Ninewells Hospital and Medical School, University of Dundee, Dundee DD1 9SY, UK; Biomedical Research Foundation, Academy of Athens, Athens 11527, Greece; Molecular and Clinical Cancer Sciences, Manchester Cancer Research Centre, Manchester Academic Health Sciences Centre, University of Manchester, Manchester M20 4GJ, UK

**Keywords:** Giant Cell Arteritis, Vasculitis, Senescence, IL-6, Temporal artery biopsy, Tocilizumab, Inflammaging

## Abstract

**Objectives:** Age is the strongest risk factor of Giant Cell Arteritis (GCA), implying a possible pathogenetic role of cellular senescence. To address this question, we applied an established senescence specific multi-marker algorithm in tissue artery biopsies (TABs) of GCA patients.

**Methods:** Seventy five positive TABs from GCA patients and 22 negative from patients with Polymyalgia Rheumatica (PMR) were retrospectively retrieved and analyzed. Senescent cells and their histologic origin were identified with specific cellular markers; IL-6 and MMP-9 were investigated as components of the senescent associated secretory phenotype (SASP) by triple co-staining. GCA or PMR artery culture supernatants were applied to primary skin fibroblasts with or without IL-6 blocking agent to explore the induction of IL-6 associated cellular senescence.

**Results:** Senescent cells were mainly present in GCA arteries at higher proportion compared to PMR (9.50% vs 2.66% respectively, p<0.0001) and were mainly originated from fibroblasts, macrophages and endothelial cells. IL-6 was expressed by senescent fibroblasts and macrophages while MMP-9 by fibroblasts only. IL-6 positive senescent cells were associated with the extension of vascular inflammation (adventitial limited disease vs transmural inflammation: 10.02% vs 4.37% respectively, p<0.0001). GCA but not PMR artery culture supernatant could induce IL-6-associated senescence that was partially inhibited by IL-6 blockade.

**Conclusions:** Senescent cells with inflammatory phenotype are present in GCA arteries and are associated with the tissue inflammatory bulk. These findings might suggest a potential implication in disease pathogenesis by perpetuating inflammation and affecting vascular remodeling via IL-6 dependent mechanisms.

## Introduction

Giant cell arteritis (GCA), is a chronic autoimmune disease, characterized by remarkable heterogeneity of the clinical phenotype and histologic pattern, reflecting the complexity of the underlying pathogenetic mechanisms which still remain unclear [1-3]. The histopathologic hallmark of the disease is the pronounced inflammation of the vascular wall, mediated by different cell types including dendritic cells, T lymphocytes, monocytes, macrophages, endothelial cells, and fibroblasts. The interplay is mediated by a variety of pro-inflammatory cytokines as well as growth and angiogenic factors that promote inflammation and abnormal tissue remodeling within the vascular wall [4]. Despite the large array of the involved cytokines, very few have been proven to play an important role in GCA [2, 5, 6] pathogenesis such as IL-6, IL-17A and GM-CSF, as attested by *in vitro* studies and the beneficial therapeutic effect of certain blockading agents and particularly the anti-IL-6 receptor antibody, tocilizumab [2, 5, 6]. However, GCA patients may still be refractory or relapse even after the introduction of novel treatments, pointing out that the major players of vascular inflammation have been only partially recognized.

GCA affects adults over 50 years old and among other risk factors, age is the strongest for GCA development [7, 8]. Vascular ageing in GCA has been associated with structural changes and altered immune responses [9], leading to the assumption that cellular senescence as part of the normal ageing of vessels, is anticipated to be involved in disease pathogenesis. Cellular senescence is a state characterized by cell cycle arrest, deregulated metabolism, macromolecular damage, and a specific secretory phenotype termed SASP (senescence associated secretory phenotype) [10]. The SASP includes pro-inflammatory cytokines, chemokines, metalloproteinases, angiogenic and growth factors implicated in wound healing, tissue plasticity and chronic inflammation. The inflammatory mediators of SASP (e.g. IL-6, IL-1β, TNF-α) may act in an autocrine and paracrine manner to induce and spread senescence [11, 12], perpetuating the inflammatory process through a local positive feedback loop. Although, the phenomenon of senescence has been well studied *in vitro* by detecting SA-β-gal, *in vivo* studies of human tissues require fresh biological material that renders the SA-β-gal approach ineffectual. On the other hand, the widely used approach to study senescence based on p16^INK4A^ and p21^WAF/Cip1^ molecular pathways has led to misleading results, since several non-senescent conditions may be linked to upregulation of these particular pathways [13, 14]. To overcome these obstacles, a multi marker algorithm for cellular senescence, has been developed and adopted by the International Cellular Senescence Association [10, 15]. This novel strategy has offered the opportunity to identify senescent cells in tissue specimens and study the pathogenetic role of cellular senescence in human diseases such as Hodgkin’s lymphoma and COVID-19 [16, 17].

So far, studies focusing on cellular senescence in GCA are limited. The microRNAs, mir-146a, mir-21 and mir-155 that have been proposed to be involved in inflammatory and senescence processes, are upregulated in TABs of GCA patients [18, 19]. In a recent study, senescence related molecular mechanisms as defined by p16^INK4A^ and p21^WAF/Cip1^ upregulation have been described in tissue specimens of GCA patients, although true senescent cells were not detected [20]. The current study aims to identify the various senescent cell types in inflamed temporal arteries of GCA and PMR patients, by applying the novel multi-marker algorithm, define the SASP key molecules and explore possible implications of senescence in GCA pathogenesis.

### Patients and Methods

Seventy five patients with GCA and 22 with PMR who were diagnosed and followed up in 3 Rheumatology centers from Greece and Italy (University of Athens, Sismanoglio General Hospital, Reggio Emilia Hospital) between 2009 and 2022, were retrospectively included in the current study. All participants underwent temporal artery biopsy as part of standard of care and fulfilled the international classification criteria for each disease [21, 22]. Active disease, either for GCA or PMR was defined by the presence of clinical symptoms and increased acute phase reactants [Erythrocyte sedimentation rate (ESR) > 20 mm/h and C-reactive protein (CRP) > 5 mg/L). The extent of vascular involvement in GCA patients as well as the absence of underlying subclinical vasculitis in PMR patients were confirmed by 18FDG Positron Emission Tomography with computed tomography (PET/CT) in almost all, except 6 patients with GCA. All diagnostic and therapeutic interventions were part of standard of care and according to physicians’ judgement. The medical records of the patient were reviewed, and all clinical and laboratory features were recorded. The study was approved by the local ethical committees of all the involved Institutions (1718016656), after obtaining patients’ informed consent and in compliance with the general data protection regulations (GDPR) of the European Union and Declaration of Helsinki. Tissue artery biopsies were stained by immunohistochemistry or immunofluorescence to detect senescent cells, their histologic origin and the components of SASP. Specific protocols were followed to quantify senescent cells per TAB. Several correlations were explored between the proportion of senescent cells in TABs and clinical or laboratory parameters. To investigate if artery culture supernatant from GCA or PMR patients has the capacity to induce IL-6 associated cellular senescence, an *ex vivo* short-term artery tissue culture system was developed to collect the 24h supernatant form GCA or PMR arteries. Subsequently, primary skin fibroblasts were treated for 5 days with the 24h temporal artery culture supernatant with or without IL-6 or IL-1β blockade, while 2 normal arteries from non-inflammatory individuals served as controls (Ctr). Bulk RNAseq analysis in already published data from TABs of GCA and non GCA patients was conducted to investigate the possible involvement of genes and pathways associated with cellular senescence. A more detailed description of methods and protocols used in the present study are provided in the supplementary material.

### Statistical analysis

For continuous variables, 2-tailed Student’s t test or Mann-Whitney were used, after implementing the Shapiro-Wilk normality test. For comparing more than two groups, one-way ANOVA with Bonferroni post hoc or Kruskal-Wallis, were performed accordingly. Associations between continuous variables were explored by Pearson’s correlation and between continuous and categorical by point-biserial correlation. All data were analyzed using GraphPad Prism v9.0 software (San Diego, California USA, www.graphpad.com). Data are presented as mean ± SD. Differences were considered statistically significant at p < 0.05.

## RESULTS

### Senescent cells are present in GCA arteries

The clinical and laboratory features of GCA and PMR patients who participated in the study are presented in **Supplementary Table S1**. In order to detect senescent cells in tissue artery biopsies (TABs) of GCA and PMR patients, the multi-marker algorithm that is wide-accepted by the international cellular senescence association was applied (ICSA) [10, 15]. Initially, single staining for GL13, an established senescence biomarker, was performed immunohistochemically in 75 GCA and 22 PMR tissue specimens, followed by immunofluorescence in 15 GCA and PMR TABs. GL13 positive senescent cells were detected in all TABs of GCA patients with both immunohistochemistry and immunofluorescence, as opposed to PMR patients, in whom senescent cells were almost absent (**Figure 1A**). In an attempt to quantify senescent cells, GL13 positive cells were calculated immunohistochemically, as a proportion of the total number of cells within TABs and after careful exclusion of the inflammatory cells in TABs of GCA patients. It was found that TABs of GCA patients (n=75) contained significantly higher proportion of senescent cells (range: 0.8%-25%) compared to PMR patients (n=22, range: 0%-6.8%) (mean proportion of GL13 positive senescent cells: 9.50% vs 2.66%, p<0.0001) (**Figure 1B**). The finding of senescent cells in GCA arteries was further confirmed, according to the second step of the algorithm, by co-staining with GL13 and p21^WAF1/Cip1^ (n=10, for both GCA and PMR group) (**Figure 1C**). Transmission electron microscopy (TEM) analysis in 2 representative samples from both groups showed that TABs from GCA patients disclosed lipofuscin accumulation (**Figure 1D**) in contrast to PMR samples (**Figure 1E**). Ultrastructural study of the arteries showed in GCA patients, cells (fibroblast) with lipofuscin granules were localized mainly near the nucleus (**Figure 1Di**) or scattered in the cytoplasm (**Figure 1Dii**). In PMR patients, cells containing lipofuscin granules were not observed (**Figure 1Ei,ii**). Interestingly, bulk RNAseq analysis of already published data from TABs of GCA and non-GCA controls, was in line with our findings showing upregulation of genes and molecular pathways linked to cellular senescence (**Supplementary Figure 1**). No correlation was found between the proportion of senescent cells in tissue specimens and the levels of CRP, ESR, the histologic pattern (transmural vs limited to adventitia), the disease phenotype (cranial vs extracranial) or the occurrence of ocular manifestations in GCA patients (**Supplementary Figure 2**).

**Figure 1.**
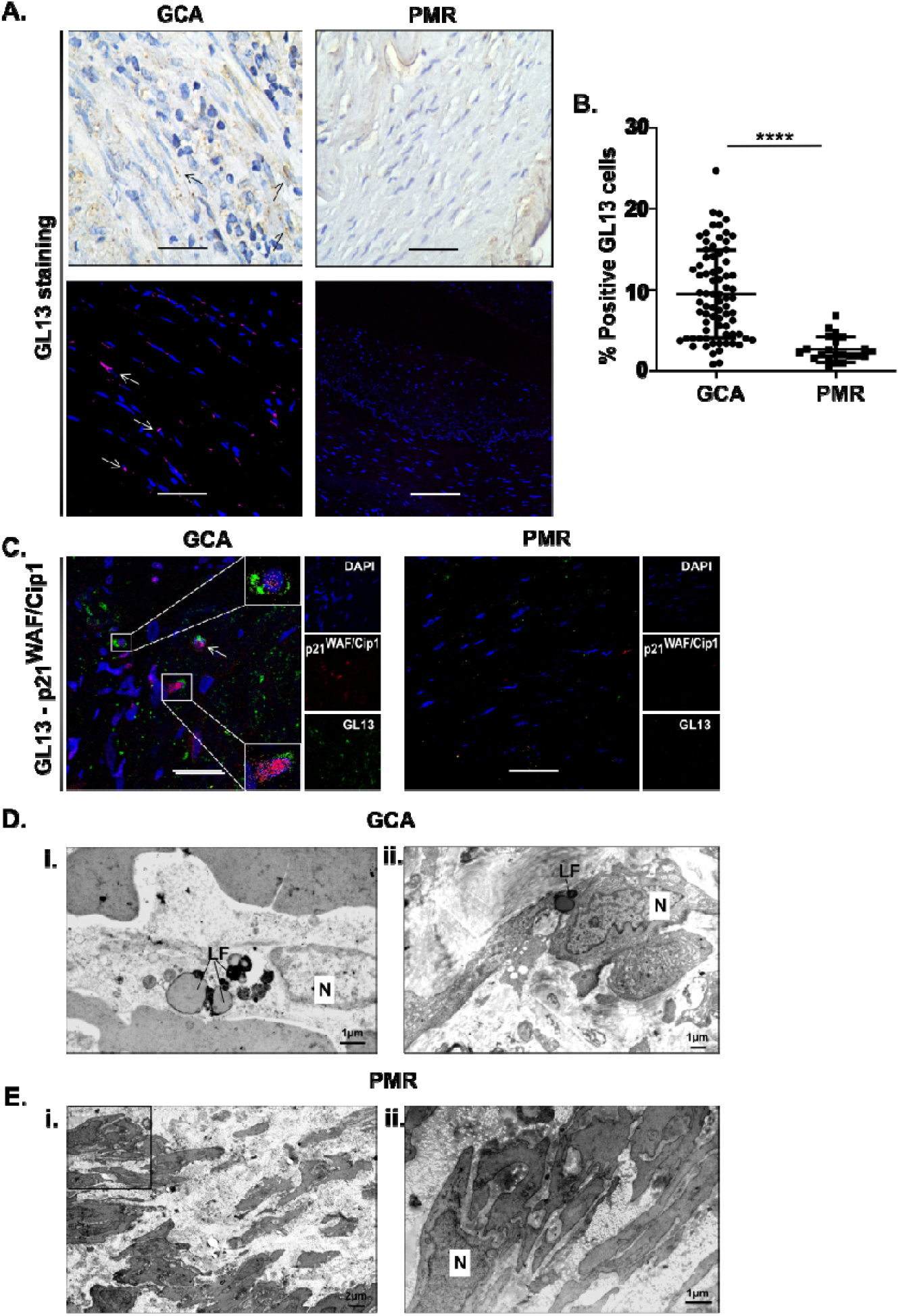
Detection of senescent cells in tissue artery biopsies of GCA patients. (A) Representative images for single GL13 staining with immunohistochemistry (upper panel) and immunofluorescence (lower panel) in TABs of GCA and PMR patients, showing higher proportion of GL13 positive cells adjacent to the inflammatory cells in GCA arteries compared to PMR. (B) Graphical representation of the proportion of GL13 positive senescent cells, after immunohistochemical evaluation by two independent readers and quantification analysis in 75 GCA and 22 PMR TABs (GCA vs PMR): 9.50% vs 2.66%, p<0.001. (C) Confirmation of senescent cells by co-staining for GL13 and p21^WAF1/Cip1^ in a TAB of a GCA compared to a PMR patient (representative image). (D) Electron micrographs of senescent cells in artery of GCA patients showing Lipofuscin (LF) granules in their cytoplasm (Di,ii). (E) Electron micrographs of cells in artery of PMR patient without lipofuscin granules (Ei,ii). Higher magnification of the area in the black box of (Ei). N: nucleus. Staining: uranyl acetate/lead citrate. Objectives (A) 20x, (C) 63x. Scale bars: 50μm (A) 10μm (C). 1μM (Di,ii, Eii), 2μM (Ei). ****p < 0.0001

### Senescent cells in GCA arteries are mainly originated by fibroblasts, macrophages and endothelial cells

Next, the origin of senescent cells was investigated by carrying out double staining with GL13 and a specific marker for different cell types each time, in TABs of 13 GCA and 13 PMR patients: (Vimentin^(+)^) for fibroblasts, (CD68^(+)^) for macrophages, (CD34^(+)^) for endothelial cells and (aSMA^(+)^) for smooth muscle cells. Double positive GL13/Vimentin (senescent fibroblasts), GL13/CD68 (senescent macrophages) and GL13/CD34 (senescent endothelial cells) cells were mainly observed in TABs of GCA patients compared to PMRs patients (**Figure 2A-C**). Additionally, double positive GL13/aSMA cells (senescent smooth cells) were hardly detected in GCA and PMR tissue samples (**Figure 2D**). The proportion of senescent cells per cell type, was calculated as the fraction of double positive cells over the total population of the specific cell type. The proportion of double positive GL13/Vimentin (senescent fibroblasts), GL13/CD68 (senescent macrophages) and GL13/CD34 (senescent endothelial) cells was higher in TABs of 13 GCA patients compared to 13 PMRs patients (29.6% vs 2.16 % p<0.001, 16.23% vs 1.7% p<0.01, and 14.53% vs 1.1% p<0.05, respectively) (**Figure 2E**). Interestingly, senescent fibroblasts in TABs of GCA patients were mainly observed in tunica media.

**Figure 2.**
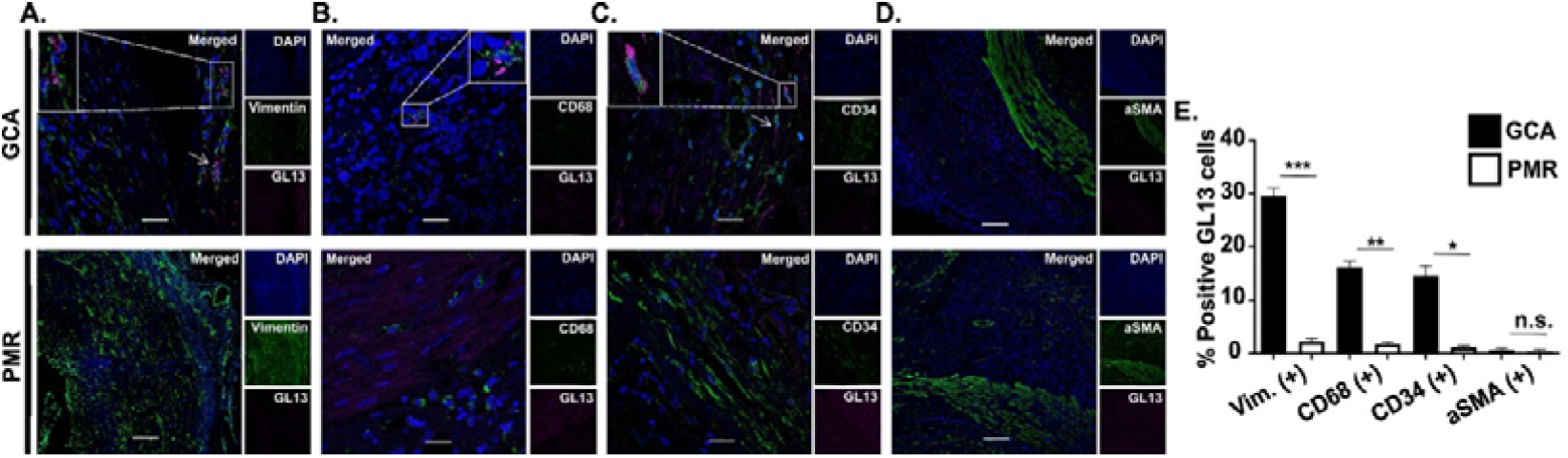
Characterization of senescent cells per cell type in tissue artery biopsies of GCA patients. Representative images of double positive senescent cells after staining for GL13 and (A) Vimentin^(+)^ (fibroblasts), (B) CD68^(+)^ (macrophages), (C) CD34^(+)^ (endothelial cells) or (D) aSMA^(+)^ (smooth cells) in a TAB of one GCA and one PMR patient (double staining immunofluorescence). Squares depict magnified cells with presence of double positive staining. (E) Quantification analysis by two independent observers, showing significantly higher proportion of senescent fibroblasts (GL13^(+)^/Vimentin^(+)^), macrophages (GL13^(+)^/CD68^(+)^) and endothelial cells (GL13^(+)^/CD34^(+)^) in 13 GCA compared to 13 PMR biopsies. Objectives 20x. Scale bars: 50 μm. n.s. not significant, *p < 0.05, **p < 0.01, p < 0.001

### Senescent cells of GCA arteries demonstrate a SASP phenotype encompassing IL-6 and MMP-9

To assess the SASP in TABs of GCA patients, triple staining in tissue specimens from 13 GCA and 13 PMR patients was applied: GL13, a specific cell type biomarker (Vimentin^(+)^, CD68^(+)^, CD34^(+)^ or aSMA^(+)^) and IL-6 which is considered a key molecule in the pathogenesis of the disease. IL-6 expression was hardly observed by double positive GL13/aSMA (senescent smooth muscle cells) cells in both GCA and PMR tissue specimens (**Figure 3A**). IL-6 was mainly expressed by double positive GL13/Vimentin positive (senescent fibroblasts) and to lesser extent GL13/CD68 (senescent macrophages) by cells in GCA arteries (**Figure 3B, C**) while double positive GL13/CD34 cells exhibited very low expression of IL-6 (**Figure 3D**). Quantification of the proportion of IL-6 positive senescent cells in TABs of 10 GCA compared to 10 PMR patients, showed higher rates of IL-6 positive cells of double positive GL13/CD68 (senescent macrophages) (7.76% vs 0.33% p<0.05 respectively) and GL13/Vimentin (senescent fibroblasts) (21.96% vs 0.73%, p<0.01 respectively) cells. No statistically significant difference was found between GCA and PMR specimens regarding the proportion of IL-6 positive cells of double positive GL13/aSMA (senescent smooth muscle cells) and GL13/CD34 (senescent endothelial cells) cells (**Figure 3E**). Given the role of metalloproteinases in tissue remodeling, the expression of MMP-9 by fibroblasts only, in 3 TABs from GCA and 3 from PMR patients, was explored. MMP-9 staining was almost exclusively observed on GL13/vimentin double positive cells of GCA arteries (**Figure 3F**) and the proportion of MMP-9 positive senescent fibroblast was higher in GCA specimens compared to PMR (13.43% vs 1.4% p<0.05) (**Figure 3G**).

**Figure 3.**
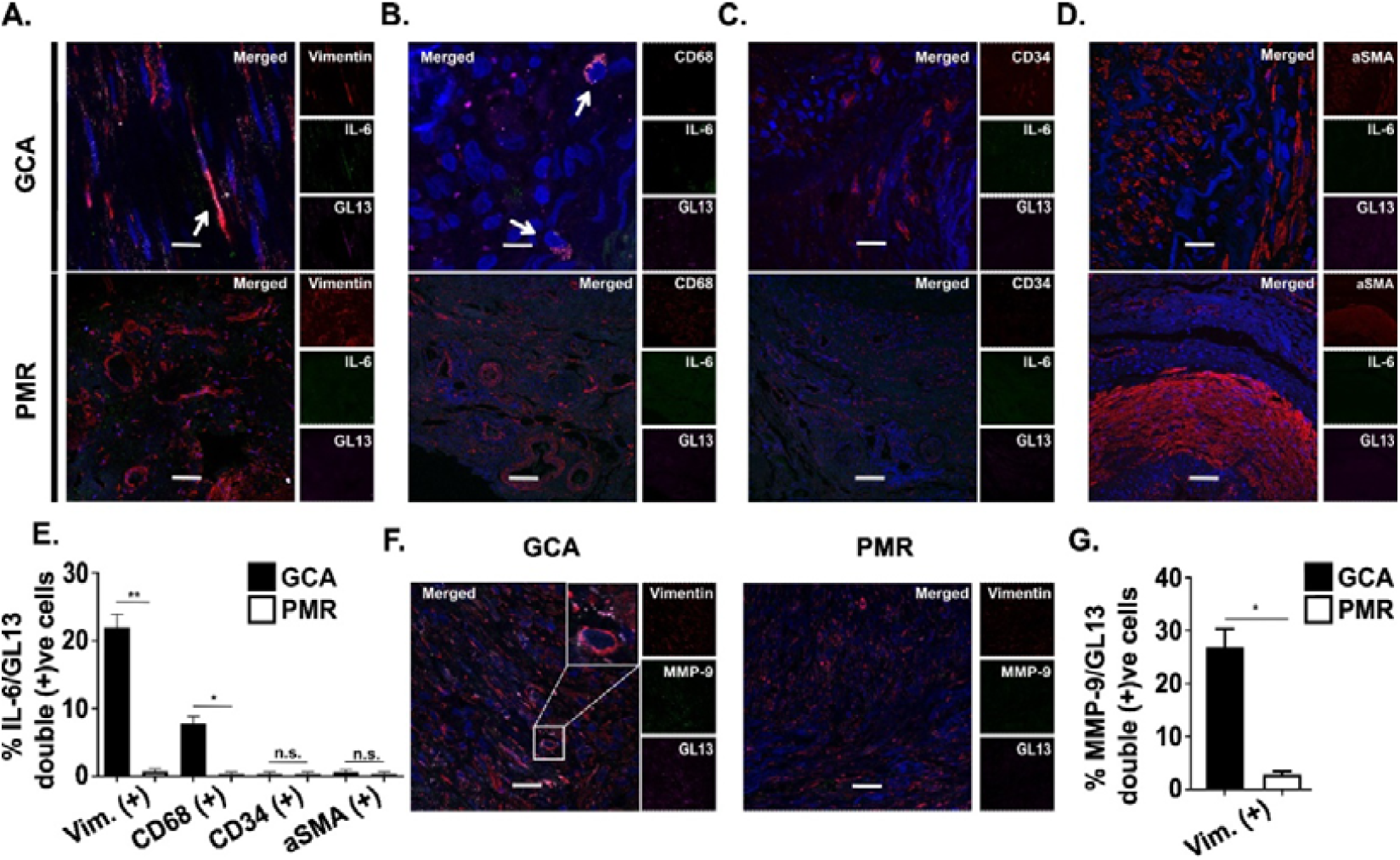
Senescent cells in GCA express SASP that includes IL-6 and MMP-9. Representative images of IL-6 expression by senescent cells of different origin: (A) GL13^(+)^/Vimentin^(+)^ (senescent fibroblasts, (B) GL13^(+)^/CD68^(+)^ (senescent macrophages), (C) GL13^(+)^/CD34^(+)^ (senescent endothelial cells)) and (D) GL13^(+)^/aSMA^(+)^ (senescent smooth muscle cells) in a TAB of n=10 GCA and n=10 PMR patients (triple staining immunofluorescence). (E) Graphical representation of the proportion of IL-6 positive senescent cells in 10 GCA and 10 PMR biopsies, showing significantly higher proportion of IL-6 positive senescent fibroblasts (GL13^(+)^/Vimentin^(+)^) and macrophages (GL13^(+)^/CD68^(+)^). Representative images of MMP-9 expression by GL13/vimentin double positive (senescent fibroblasts) cells in a TAB of one GCA and one PMR patient (triple staining immunofluorescence). (F-G) Graphical representation showing higher proportion of MMP-9 positive senescent fibroblast (GL13^(+)^/vimentin^(+)^) in the 3 GCA compared to 3 PMR specimens. Objectives (A), (D) 20x, (B), (C), 63x. Scale bars: (A), (D) 50 μm, (B), (C), 10μm. n.s. not significant, *p < 0.05, **p < 0.01

### The presence of IL-6 positive senescent cells is associated with the extent of inflammation in GCA arteries

To explore the association of IL-6 associated senescent cells with the extent of the inflammatory process in GCA arteries, we compared 7 patients with disease limited to the adventitia (limited disease) and 68 patients with transmural inflammation (extended disease) (**Figure 4A**). GL13/IL-6 double positive cells were more prominent in GCA patients with extended disease compared to those with limited disease (**Figure 4B**) and after quantification the proportion of IL6+ senescent cells was found significantly higher in the former group (10.02% vs 4.37%, p<0.0001)(**Figure 4C**).

**Figure 4.**
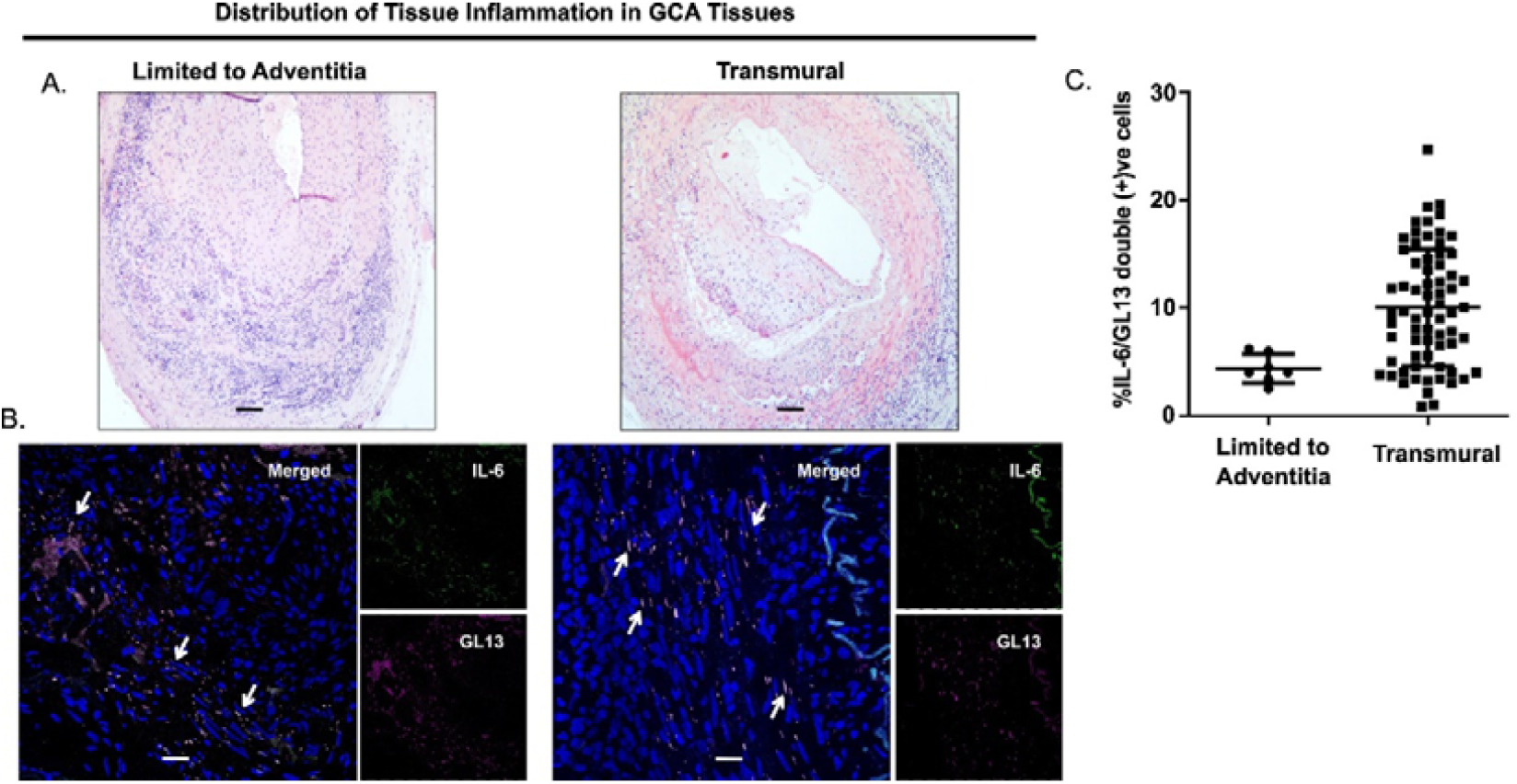
IL-6 positive senescent cells in GCA biopsies of patients with limited or extended disease. (A) Representative images of TAB from GCA patient with disease limited to adventitia (left panel) and with extended transmural inflammation (right panel) (H&E). (B) Double positive IL-6/GL13 senescent cells are observed more abundantly in extended disease (right panel) compared to limited disease (representative images from one GCA patient from each group, immunofluorescence). (C) Graphical representation of the proportion of IL-6 positive senescent cells in 7 GCA patients with limited vs 68 patients with extended disease. Positive staining for each panel was quantified by two independent observers. Objectives (A) 10x, (B) 20x. Scale bars: (A) 100μm, (B) 50μm. ****p < 0.0001

### GCA artery culture supernatant triggers senescence phenotype in primary fibroblasts. Significant inhibition by IL-6 signaling blockade

To investigate if the microenvironment of GCA arteries carries the capacity to induce IL-6 associated cellular senescence, we treated primary skin fibroblasts for 5 days with 24h temporal artery culture supernatant from 3 patients with GCA and 3 with PMR (disease controls) while 2 normal arteries from healthy individuals served as normal controls (Ctr). IL-6 positive senescent fibroblasts were detected by double staining for GL13 and IL-6. Inhibition experiments were also performed by applying IL-6 or IL-1β signaling blocking agents (**Figure 5A**). A series of preliminary experiments were performed to define the optimal settings and the appropriate negative controls (**Supplementary Figure 3**). Time dependent preliminary experiments in which primary skin fibroblasts were treated with aorta culture supernatant from one GCA patient with aortitis, showed that the optimal time to detect cellular senescence, is after 5 days of treatment (**Supplementary figure 3**). Interestingly, primary skin fibroblasts treated with sera from GCA or PMR patients for 5 days displayed no cellular senescence (data not shown). Double positive GL13/IL-6 primary skin fibroblasts were mainly observed after treatment with GCA supernatants while cellular senescence was minimal with PMR supernatants. No GL13/IL-6 positive fibroblasts were observed with supernatants from control arteries (**Figure 5B**). The proportion of double positive GL13/IL-6 fibroblasts after treatment with GCA supernatants was significantly higher compared to PMR supernatants (25.17% vs 1.50%, p<0.0001) (**Figure 5C**). Interestingly, the 24h temporal artery culture supernatant from GCA and PMR had comparable levels of IL-6 (**Supplementary Figure 4**). Next, we blocked IL-6 and IL-1β signaling, using 1μg/ml tocilizumab and 10 μg/ml anakinra respectively as described previously [23, 24], before treating primary skin fibroblasts with GCA temporal artery culture supernatants (**Figure 5D**). Induction of IL-6 associated cellular senescence was significantly inhibited by IL-6 receptor blockade (12.00% vs 32.17%, p < 0.001) and to a lesser extent by IL-1β receptor inhibition (25.37% vs 32.17%, p < 0.05) (**Figure 5D**). IL-6 and IL-1 receptor blockade, inhibited IL-6 associated cellular senescence by approximately 60% and 10%, respectively. Moreover, the inhibition of IL-6 associated cellular senescence observed by IL-6 receptor blocking was statistically higher compared to IL-1 blocking (12.00% vs 25.37%, p < 0.01) (**Figure 5E**).

**Figure 5.**
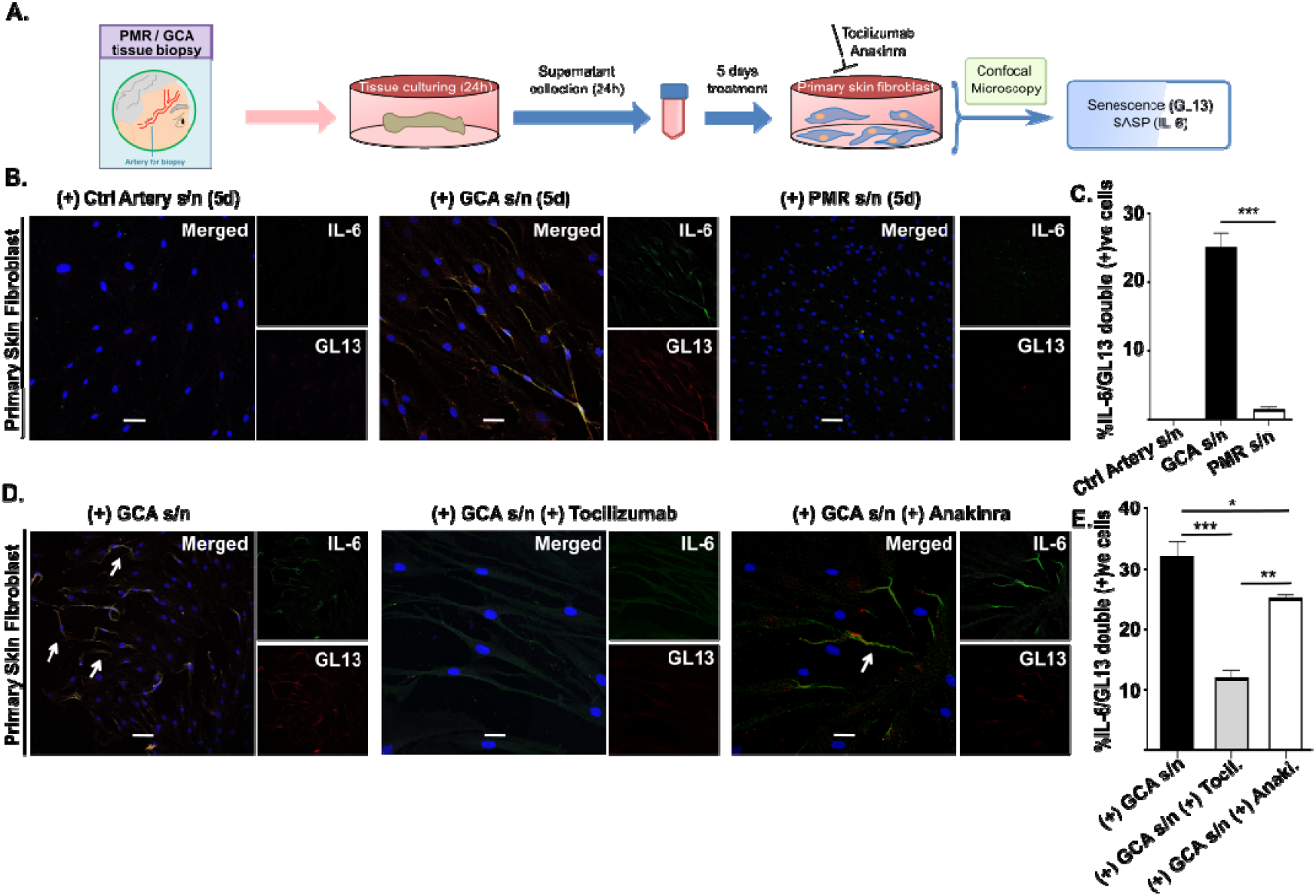
IL-6 associated cellular senescence induced by GCA temporal artery culture supernatant and partial inhibition by IL-6 blockade. (A) Workflow of experimental procedure. (B) Representative images from IL-6/GL13 co-staining of primary skin fibroblasts treated with GCA (left panel), PMR (middle panel) and healthy artery (right panel) tissue culture supernatants for 5 days, respectively, demonstrating IL6-positive associated senescent fibroblasts mainly after treatment with GCA supernatant. (C) Graphical representation of the percentage of double positive IL-6/GL13 fibroblasts after treatment with GCA (n=3), PMR (n=3) or ctr supernatants (n=2). (D) Primary skin fibroblasts were pre-treated with IL-6 or IL-1β receptor antagonists for 2 hours before treatment with GCA tissue culture supernatant for 5 days. Representative images from IL-6/GL13 co-staining of primary fibroblasts treated with GCA tissue culture supernatant only right panel (left panel), GCA tissue culture supernatant plus tocilizumab (IL-6 receptor blocking agent) (middle panel) or GCA tissue culture supernatant plus anakinra (IL-1β receptor blocking agent) (right panel). Tocilizumab resulted in partial inhibition of IL-6 associated senescence of treated fibroblasts and anakinra to a lesser extent. (D) Graphical representation of the percentage of IL-6/GL13 double positive senescent fibroblasts in (C), showing that tocilizumab has the most significant inhibitory capacity (E). Positive events for each staining were quantified by two independent observers). Objective 63 x. Scale bar: 20 μm. *p < 0.05, **p <0.01, ***p < 0.001

## Discussion

Our study presents some unique features regarding cellular senescence in GCA. To the best of our knowledge, this is the first study that true senescent cells are directly identified in the diseased tissues of a non-neoplastic, inflammatory autoimmune disease, following an established multi-marker algorithm. The latter was also applied recently by our team in a COVID-19 cohort unrevealing the role of SARS-CoV2 in triggering cellular senescence [16]. This particular approach represents a novelty in the field of systemic inflammatory diseases, as opposed to the usual strategy of applying solely p16^INK4A^ and/or p21^WAF/Cip1^ staining for the identification of senescence, leading in many occasions to the misinterpretation of the latter [10, 14, 15, 20, 25]. On the contrary, by employing the multi-marker approach, true senescent cells can be strictly identified, offering a unique scientific “tool” to understand and evaluate important biologic properties such as the exact location and quantitative burden of senescent cells, their origin in terms of cell type and their SASP components that define their pathogenetic potential [10, 15]. The application of the multi-marker algorithm in the present study, led to significant and novel findings. First, we showed that although senescent cells could be also detected in age and sex matched PMR TABs-most likely as a result of “normal” vessel ageing-their proportion is significantly increased in GCA TABs. The lack of inflammatory infiltrate in PMR TABs, suggests an association between the inflammatory process within the arteries and the occurrence of senescent cells, supporting the notion that in GCA arteries there are both pre-existing and disease associated senescent cells. Indeed, senescent cells in TABs of GCA patients are mainly observed in close proximity with the inflammatory infiltrate. Second, it was demonstrated that cellular senescence in GCA arteries is limited to specific cell populations, most prominently in fibroblasts, and macrophages and to a lesser extent endothelial cells. In this way, cellular senescence can be conceptualized through specific cell types in terms of histologic features, function and location within the layers of the vessel. Third, the triple staining for SASP components, driven by the origin of senescent cells and already known pathogenetically important molecules, revealed that IL-6 and MMP-9 positive senescent cells are present in GCA arteries, confirming that SASP in GCA carries an inflammatory and impaired vascular remodeling potential. Fourth, by quantifying IL-6+ senescent cells, it was shown that the IL6+ senescent burden is proportional to the histologic pattern/extension of the disease, underlining an important link among IL-6, degree of inflammation and cellular senescence in GCA. Finally, and apart from the studies at the biopsy levels, functional experiments demonstrated that the microenvironment of GCA has the capacity to induce, through soluble mediators including IL-6, IL-6 associated cellular senescence, clearly implying a positive IL-6 dependent loop between the disease itself and cellular senescence.

Our findings provide important insights in GCA pathogenesis and place cellular senescence as a potential contributor of the inflammatory process within the diseased arteries. Senescent cells probably pre-exist in temporal arteries of GCA patients as supported by our findings in the TABs of age and sex matched PMR patients, reflecting the “normal” background ageing burden of cellular senescence. Apart from the bulk of pre-existing senescent cells, an excess of senescent cells was documented within the GCA arteries in our study, that most likely is related to the inflammatory burden of the disease and represents inducible senescent cells. Whether GCA arteries display higher degree of pre-existing senescent cells or are more vulnerable to cellular senescence is difficult to be dissected, although recent studies have shown that environmental stimuli may provoke premature senescence state [26]; nevertheless our work clearly showed that soluble factors derived from the microenvironment of GCA and not PMR arteries, has the capacity to induce senescence and that IL-6 plays a significant role as attested by the inhibition experiments of IL-6 blockade. The fact that IL-6 blockade resulted in significant but not total inhibition of cellular senescence, suggests that other factors such as GM-CSF or IL-17A may act synergistically to induce the senescence phenomenon. Thus, the inflamed temporal arteries during disease course are thought to release soluble mediators, with IL-6 playing a major role, that generate and increase the burden of senescent cells within the vessel.

The excess of senescent cells in GCA arteries is of great clinical significance with important biologic consequences, considering their unique features. Despite the fact that senescent cells are characterized by cell cycle arrest with loss of proliferative capacity, they are resistant to apoptosis and cannot be easily removed and cleared from tissues [10, 15]. The presence and persistence of senescent cells has biologic effects which are largely defined by two factors: i) the histologic origin of senescent cells [27-29] and ii) the SASP components. The acquired properties of senescent cells contribute to their dynamic functional state and renders them capable of interfering with the neighboring structures. In this direction, we found that senescent cells originated mainly by fibroblasts and macrophages in GCA arteries, express key molecules such as IL-6 and MMP-9. The IL-6 derived from senescent cells may act in an autocrine and paracrine manner either to induce and spread cellular senescence to non-senescent neighboring cells or to amplify the local inflammatory milieu [10, 30]. Interestingly, it was recently shown that chronic stimulation of macrophages through TLR2 signaling may drive granuloma formation, a histologic feature of GCA and simultaneously DNA damage response, a well-known trait of cellular senescence [31]. Similarly, MMP9 may contribute to the delay of the resolution of inflammation and deregulation of vascular remodeling which are closely related to aneurysm formation and refractoriness to treatment in GCA [32]. The role of endothelial senescent cells is less obvious but previous data have shown that cellular senescence is associated with overexpression of ICAM-1 by the endothelium, rendering them more inflammatory phenotype [33]. Overall, it appears that the pathogenetic model of GCA could be enriched by integrating cellular senesce in a similar way to that observed in severe COVID-19 infection [10, 34]. According to this model, pre-existing and inducible senescent cells in GCA arteries may play an active role in pathogenesis through perpetuation and maintenance of the inflammatory process as well as by interfering adversely with the tissue repair and remodeling of the inflamed vessel. In this way, significant co-morbidities of GCA and mainly atherosclerosis can be further interpreted by the interference of foamy senescent macrophages which have been proposed to display atherogenic functions [27, 29]. This model is further supported by the fact that the proportion of senescent cells is correlated with the extent of inflammation within the affected arteries. In this line, the RNAseq and gene ontology analysis of already published data from GCA arteries performed by us, revealed an enrichment of genes and molecular pathways related to cellular senescence. Furthermore, GWAS in GCA, pointed out specific SNP of important SASP components such as IL-6 or other cytokines, that may confer genetic predisposition to GCA through cellular senescence [35]. Finally, the epigenetic regulation of cellular senescence has been recognized to be involved in pathogenesis of cardiovascular diseases [36] and especially in GCA specific microRNAs linked to senescence were found upregulated in GCA arteries [18].

Undoubtably, the application of the multi-marker algorithm is one of the major strengths of this study. This approach was crucial to draw objective conclusions and “quantify” cellular senescence, opening a new avenue to study the role of senescence in tissue samples of GCA. Another strength is the large number of TABs that were studied and analyzed by the same scientific team to ensure statistical power, accuracy and reproducibility of the research findings. One important weakness is that there was a limited investigation of SASP components, including only IL-6 and MMP-9. It is more than certain that SASP, as a route of pathogenicity, is characterized by significant diversity that incorporates among others IL-17A and GM-CSF, and therefore these molecules should be extensively investigated in future studies. Another limitation is the use of glucocorticosteroids (GCS) by a significant number of patients prior to TAB. Although the inflammatory milieu in TAB has been proved not to be affected by short-term course of GCs [37], their impact on senescence induction in unknown. However, in our study the proportion of senescent cells among GCA patients who have received GCS for different duration ranging from 1 to > 30 days, was not statistically different compared to untreated GCA patients. The lack of TABs from GCA patients in remission is also a weakness of our study. A second TAB after treatment administration would offer the opportunity to study the biologic behavior and features of senescent cells in the inactive state of the disease. Finally, additional functional experiments are required to identify other than IL-6 molecules that could mediate cellular senesce and have a role in GCA pathogenesis.

In conclusion, senescent cells are present in GCA arteries and display an IL-6/MMP-9 SASP that mediates inflammation and impaired vascular remodeling. Especially, IL6+ senescent cells follow the extension and bulk of the inflammatory infiltrate and can be induced by soluble mediators of the microenvironment of GCA arteries in an IL-6 dependent way, through an IL-6 centered positive loop. The plasticity and diversity of senescent cells ensure their active participation in GCA pathogenesis, and their detection in tissue biopsy may serve as a potential tissue biomarker for disease severity, response to treatment and prognosis of serious complications. In addition, the implication of cellular senescence in GCA pathogenesis and the IL-6 dependent vicious cycle of inflammation, sets reasonable questions on the efficacy of senolytic treatment as a complementary strategy in the management of GCA. Future studies focusing on these issues are anticipated to shed light at both clinical and molecular level regarding the pathogenesis, diagnosis and treatment of GCA.

## References

1. Ciccia F, Macaluso F, Mauro D, Nicoletti GF, Croci S, Salvarani C. New insights into the pathogenesis of giant cell arteritis: are they relevant for precision medicine? The Lancet Rheumatology. 2021; 3(12):e874–885.

2. Stone JH, Spotswood H, Unizony SH, Aringer M, Blockmans D, Brouwer E, et al. New-onset versus relapsing giant cell arteritis treated with tocilizumab: 3-year results from a randomized controlled trial and extension. Rheumatology (Oxford). 2022 Jul 6; 61(7):2915–2922.

3. van der Geest KSM, Sandovici M, van Sleen Y, Sanders JS, Bos NA, Abdulahad WH, et al. Review: What Is the Current Evidence for Disease Subsets in Giant Cell Arteritis? Arthritis Rheumatol. 2018 Sep; 70(9):1366–1376.

4. Weyand CM, Goronzy JJ. Immune mechanisms in medium and large-vessel vasculitis. Nat Rev Rheumatol. 2013 Dec; 9(12):731–740.

5. Cid MC, Unizony SH, Blockmans D, Brouwer E, Dagna L, Dasgupta B, et al. Efficacy and safety of mavrilimumab in giant cell arteritis: a phase 2, randomised, double-blind, placebo-controlled trial. Ann Rheum Dis. 2022 May; 81(5):653–661.

6. Palamidas DA, Argyropoulou OD, Georgantzoglou N, Karatza E, Xingi E, Kapsogeorgou EK, et al. Neutrophil extracellular traps in giant cell arteritis biopsies: presentation, localization and co-expression with inflammatory cytokines. Rheumatology (Oxford). 2022 Apr 11; 61(4):1639–1644.

7. Gonzalez-Gay MA, Vazquez-Rodriguez TR, Lopez-Diaz MJ, Miranda-Filloy JA, Gonzalez-Juanatey C, Martin J, et al. Epidemiology of giant cell arteritis and polymyalgia rheumatica. Arthritis Rheum. 2009 Oct 15; 61(10):1454–1461.

8. Larsson K, Mellstrom D, Nordborg E, Oden A, Nordborg E. Early menopause, low body mass index, and smoking are independent risk factors for developing giant cell arteritis. Ann Rheum Dis. 2006 Apr; 65(4):529–532.

9. Mohan SV, Liao YJ, Kim JW, Goronzy JJ, Weyand CM. Giant cell arteritis: immune and vascular aging as disease risk factors. Arthritis Res Ther. 2011 Aug 2; 13(4):231.

10. Gorgoulis V, Adams PD, Alimonti A, Bennett DC, Bischof O, Bishop C, et al. Cellular Senescence: Defining a Path Forward. Cell. 2019 Oct 31; 179(4):813–827.

11. Birch J, Gil J. Senescence and the SASP: many therapeutic avenues. Genes Dev. 2020 Dec 1; 34(23-24):1565–1576.

12. Coppe JP, Desprez PY, Krtolica A, Campisi J. The senescence-associated secretory phenotype: the dark side of tumor suppression. Annu Rev Pathol. 2010; 5:99–118.

13. Hernandez-Segura A, de Jong TV, Melov S, Guryev V, Campisi J, Demaria M. Unmasking Transcriptional Heterogeneity in Senescent Cells. Curr Biol. 2017 Sep 11; 27(17):2652–2660 e2654.

14. Karimian A, Ahmadi Y, Yousefi B. Multiple functions of p21 in cell cycle, apoptosis and transcriptional regulation after DNA damage. DNA Repair (Amst). 2016 Jun; 42:63–71.

15. Kohli J, Wang B, Brandenburg SM, Basisty N, Evangelou K, Varela-Eirin M, et al. Algorithmic assessment of cellular senescence in experimental and clinical specimens. Nat Protoc. 2021 May; 16(5):2471–2498.

16. Evangelou K, Veroutis D, Paschalaki K, Foukas PG, Lagopati N, Dimitriou M, et al. Pulmonary infection by SARS-CoV-2 induces senescence accompanied by an inflammatory phenotype in severe COVID-19: possible implications for viral mutagenesis. Eur Respir J. 2022 Aug; 60(2).

17. Myrianthopoulos V, Evangelou K, Vasileiou PVS, Cooks T, Vassilakopoulos TP, Pangalis GA, et al. Senescence and senotherapeutics: a new field in cancer therapy. Pharmacol Ther. 2019 Jan; 193:31–49.

18. Croci S, Zerbini A, Boiardi L, Muratore F, Bisagni A, Nicoli D, et al. MicroRNA markers of inflammation and remodelling in temporal arteries from patients with giant cell arteritis. Ann Rheum Dis. 2016 Aug; 75(8):1527–1533.

19. Olivieri F, Rippo MR, Monsurro V, Salvioli S, Capri M, Procopio AD, et al. MicroRNAs linking inflamm-aging, cellular senescence and cancer. Ageing Res Rev. 2013 Sep; 12(4):1056–1068.

20. Jiemy WF, van Sleen Y, Graver JC, Pringle S, Brouwer E, van der Geest K, et al. Indication of activated senescence pathways in the temporal arteries of patients with giant cell arteritis. Arthritis Rheumatol. 2023 Apr 14.

21. Dasgupta B, Cimmino MA, Kremers HM, Schmidt WA, Schirmer M, Salvarani C, et al. 2012 Provisional classification criteria for polymyalgia rheumatica: a European League Against Rheumatism/American College of Rheumatology collaborative initiative. Arthritis Rheum. 2012 Apr; 64(4):943–954.

22. Hunder GG, Bloch DA, Michel BA, Stevens MB, Arend WP, Calabrese LH, et al. The American College of Rheumatology 1990 criteria for the classification of giant cell arteritis. Arthritis Rheum. 1990 Aug; 33(8):1122–1128.

23. Al-Jomah N, Al-Mohanna FH, Aboussekhra A. Tocilizumab suppresses the procarcinogenic effects of breast cancer-associated fibroblasts through inhibition of the STAT3/AUF1 pathway. Carcinogenesis. 2021 Dec 31; 42(12):1439–1448.

24. Nicolas AM, Pesic M, Engel E, Ziegler PK, Diefenhardt M, Kennel KB, et al. Inflammatory fibroblasts mediate resistance to neoadjuvant therapy in rectal cancer. Cancer Cell. 2022 Feb 14; 40(2):168–184 e113.

25. Alcorta DA, Xiong Y, Phelps D, Hannon G, Beach D, Barrett JC. Involvement of the cyclin-dependent kinase inhibitor p16 (INK4a) in replicative senescence of normal human fibroblasts. Proc Natl Acad Sci U S A. 1996 Nov 26; 93(24):13742–13747.

26. Liakou E, Mavrogonatou E, Pratsinis H, Rizou S, Evangelou K, Panagiotou PN, et al. Ionizing radiation-mediated premature senescence and paracrine interactions with cancer cells enhance the expression of syndecan 1 in human breast stromal fibroblasts: the role of TGF-beta. Aging (Albany NY). 2016 Aug; 8(8):1650–1669.

27. Childs BG, Baker DJ, Wijshake T, Conover CA, Campisi J, van Deursen JM. Senescent intimal foam cells are deleterious at all stages of atherosclerosis. Science. 2016 Oct 28; 354(6311):472–477.

28. Ciccia F, Rizzo A, Ferrante A, Guggino G, Croci S, Cavazza A, et al. New insights into the pathogenesis of giant cell arteritis. Autoimmun Rev. 2017 Jul; 16(7):675–683.

29. Matacchione G, Perugini J, Di Mercurio E, Sabbatinelli J, Prattichizzo F, Senzacqua M, et al. Senescent macrophages in the human adipose tissue as a source of inflammaging. Geroscience. 2022 Aug; 44(4):1941–1960.

30. Hernandez-Rodriguez J, Segarra M, Vilardell C, Sanchez M, Garcia-Martinez A, Esteban MJ, et al. Tissue production of pro-inflammatory cytokines (IL-1beta, TNFalpha and IL-6) correlates with the intensity of the systemic inflammatory response and with corticosteroid requirements in giant-cell arteritis. Rheumatology (Oxford). 2004 Mar; 43(3):294–301.

31. Herrtwich L, Nanda I, Evangelou K, Nikolova T, Horn V, Sagar, et al. DNA Damage Signaling Instructs Polyploid Macrophage Fate in Granulomas. Cell. 2016 Nov 17; 167(5):1264–1280 e1218.

32. Watanabe R, Maeda T, Zhang H, Berry GJ, Zeisbrich M, Brockett R, et al. MMP (Matrix Metalloprotease)-9-Producing Monocytes Enable T Cells to Invade the Vessel Wall and Cause Vasculitis. Circ Res. 2018 Aug 31; 123(6):700–715.

33. Gorgoulis VG, Zacharatos P, Kotsinas A, Kletsas D, Mariatos G, Zoumpourlis V, et al. p53 activates ICAM-1 (CD54) expression in an NF-kappaB-independent manner. EMBO J. 2003 Apr 1; 22(7):1567–1578.

34. Schmitt CA, Tchkonia T, Niedernhofer LJ, Robbins PD, Kirkland JL, Lee S. COVID-19 and cellular senescence. Nat Rev Immunol. 2023 Apr; 23(4):251–263.

35. Carmona FD, Vaglio A, Mackie SL, Hernandez-Rodriguez J, Monach PA, Castaneda S, et al. A Genome-wide Association Study Identifies Risk Alleles in Plasminogen and P4HA2 Associated with Giant Cell Arteritis. Am J Hum Genet. 2017 Jan 5; 100(1):64–74.

36. Paschalaki KE, Starke RD, Hu Y, Mercado N, Margariti A, Gorgoulis VG, et al. Dysfunction of endothelial progenitor cells from smokers and chronic obstructive pulmonary disease patients due to increased DNA damage and senescence. Stem Cells. 2013 Dec; 31(12):2813–2826.

37. Narvaez J, Bernad B, Roig-Vilaseca D, Garcia-Gomez C, Gomez-Vaquero C, Juanola X, et al. Influence of previous corticosteroid therapy on temporal artery biopsy yield in giant cell arteritis. Semin Arthritis Rheum. 2007 Aug; 37(1):13–19.

