## Supplemental Information for "Senescent cells in Giant Cell Arteritis have inflammatory phenotype participating in tissue injury via IL-6 dependent pathways"

|  | *GCA (n=70)* | *PMR (n=22)* |
| --- | --- | --- |
| Age (mean±SD, years) | 74.5±7.3 | 72.5±9.7 |
| Gender (female %) (n) | 65.7(46) | 63.6(14) |
| Disease course (%) (n)  Newly diagnosed  Relapse | 94.3(66)  5.7(4) | 72.7(16)  27.3(6) |
| TAB Histologic Pattern (%) (n)  Transmural  Limited to adventitia  Vasa vasoritis  Other | 82.9(58/70)  12.9(9/70)  1.4(1/70)  2.9(2/70) | - |
| Vascular Disease Extent (%) (n)  Cranial  Generalized | 60.9(39/64)  39.1(25/64) | - |
| Clinical Characteristics (%) (n)  Constitutional symptoms  Fever  Fatigue/Malaise  Anorexia  Weight loss  Cranial  Headache  Scalp tenderness  Jaw claudication  Temporal artery abnormalities  Tenderness  Decreased pulsation  Both  Musculoskeletal involvement  Arthralgia  Myalgia  Morning stiffness  Polymyalgia Rheumatica  Ocular complications  Amaurosis fugax  Blurred vision  Blindness  Diplopia  Neurologic involvement  Cranial neuropathy  Mononeuritis multiplex | 42.9(30/70)  41.4(29/70)  17.1(12/70)  12.9(9/70)  77.1(54/70)  31.4(22/70)  41.4(29/70)  13.4(9/67)  16.4(11/67)  6(4/67)  41.4(29/70)  37.1(26/70)  34.3(24/70)  45.7(32/70)  8.6(6/70)  12.9(9/70)  18.6(13/70)  11.4(8/70)  2.9(2/70)  1.4(1/70) | 18.2(4/22)  22.7(5/22)  0(0/22)  4.5(1/22)  13.6(3/22)  0(0/22)  4.5(1/22)  0(0/22)  0(0/22)  0(0/22)  95.5(21/22)  59.1(13/22)  63.6(14/22)  100(22/22)  0(0/22)  0(0/22)  0(0/22)  0(0/22)  0(0/22)  0(0/22) |
| Laboratory features  ESR (>20 mm/h) (%) (n)  ESR (mean±SD, mm/h)  CRP (>5 mg/L) (n)  CRP (mean±SD, mg/L) | 96.8(61/63)  78.4±30.9  100(68/68)  78.1±57(68) | 95.5(21/22)  70.7±33.2  86.4(19/22)  42±49.9 |
| Disease related treatment (%) (n)  Glucocorticosteroids  < 7 days  7-14 days  15-30 days  > 30 days  Immunosuppressives  Methotrexate (110-180 days)  Tocilizumab (24 months) | 61.4(43/70)  20(14/70)  10(7/70)  18.6(13/70)  12.9(9/70)  2.9(2/70)  1.4(1/70) | 36.4(8/22)  0(0/22)  13.6(3/22)  9.1(2/22)  13.6(3/22)  4.5(1/22)  0(0/22) |

**Supplementary Table 1. Baseline characteristics of GCA and PMR patients at the time tissue artery biopsy**

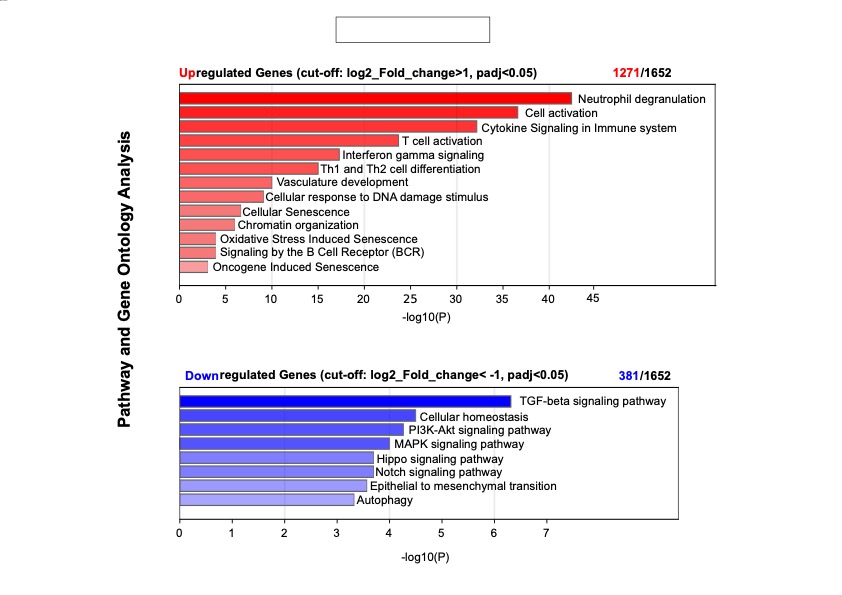

**Supplementary Figure 1. Bar graph representing selection of enriched pathways identified in TABs from GCA and non-GCA control patients.** Bulk RNA-seq analysis was performed in 20 GCA TABs sections and was compared and normalized with 20 non-GCA control TBAs. 1652 significant deregulated genes were identified, where 1271 and 381 were up- and down-regulated respectively. Gene ontology and pathway analysis was performed in both groups of genes. Pathways and biological processes with p-value lower than 0.05 were considered to be significantly enriched. X-axis depicts –log10 P-value.

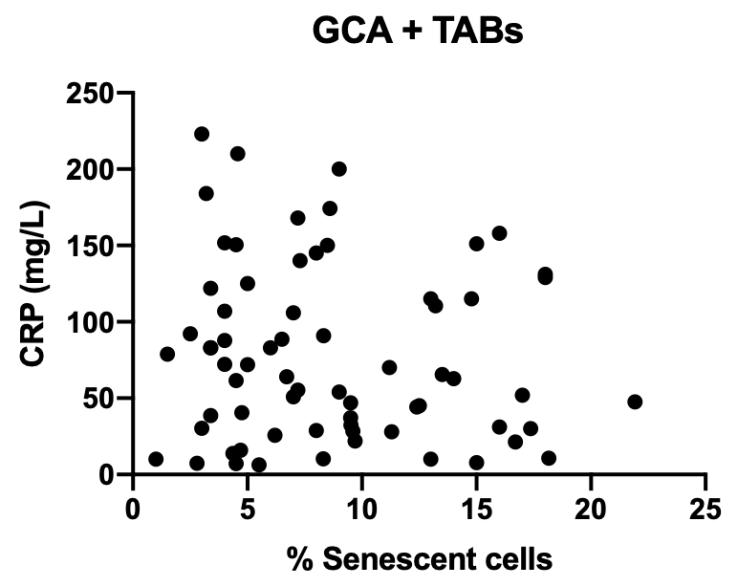

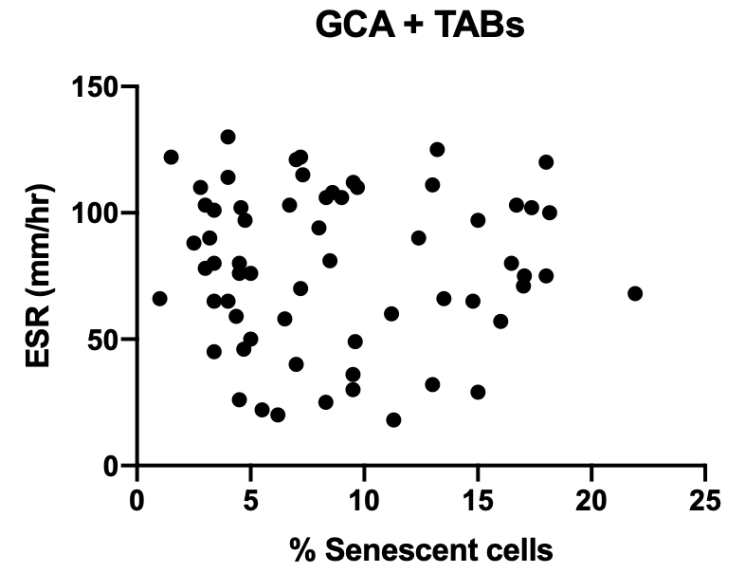

**Supplementary Figure 2. Correlations between the proportion of GL13 positive senescent cells in GCA arteries and CRP or ESR.** No significant correlation was found between the proportion of GL13 positive senescent cells and CRP (mg/l) in GCA patients (n=68)(Pearson’s correlation, r=-0.10, p=0.387) (A). No significant correlation was found between the proportion of GL13 positive senescent cells and ESR (mm/h) in GCA patients (n=63) (Pearson’s correlation, r=-0.009, p=0.943) (B).

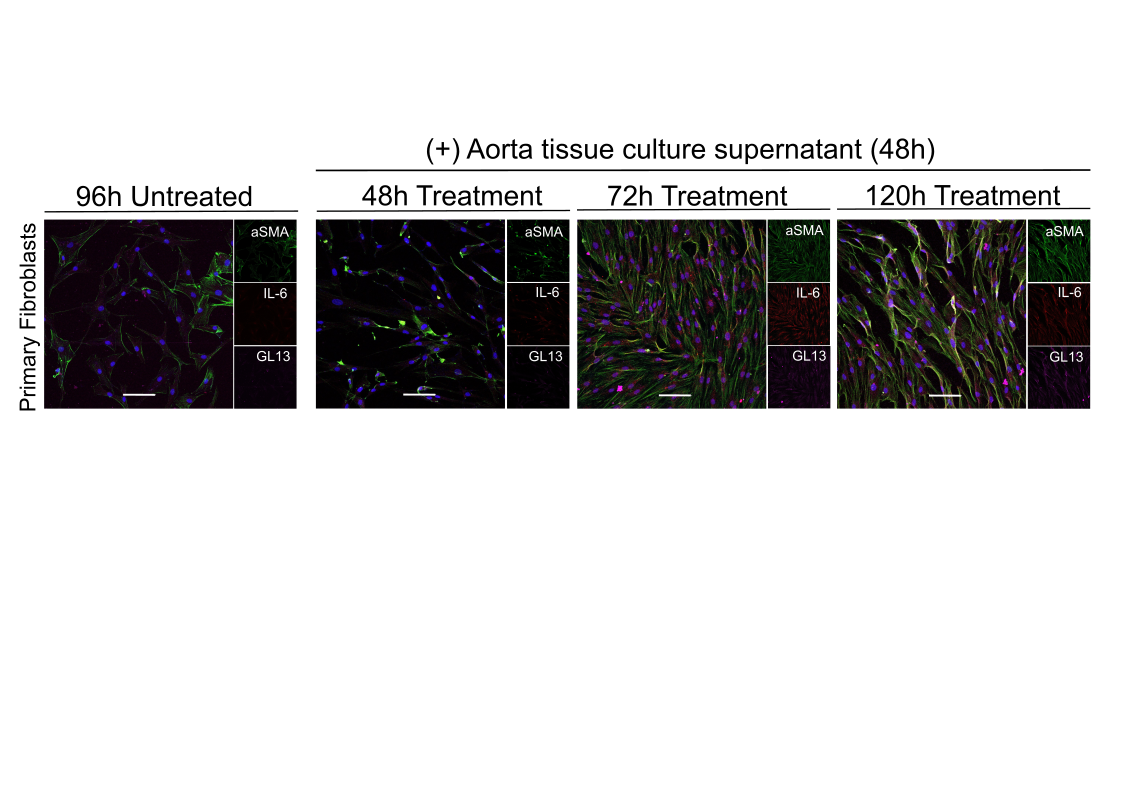

**Supplementary Figure 3. Time dependent experiments with aorta culture supernatant to determine the optimal time point for inducing IL-6 associated senescence on primary skin fibroblasts.** Aorta segment from 1 GCA patients with aortitis was cultured for 48h and the supernatant was applied 50% diluted in primary skin fibroblast for 48h, 72h and 120h. Treated and non-treated cells were fixed and stained for the detection of double positive senescence (GL13) and (IL-6) cells in the respective time points. Cellular senescence was starting observed after 72h of treatment and high level of senescent cells was observed after 120h of treatment.

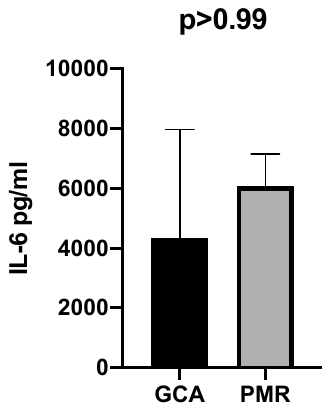

**Supplementary Figure 4. IL-6 levels of GCA and PMR temporal artery culture supernatants after 24h.** IL-6 was detected in temporal artery culture supernatants from 3 GCA and 3 PMR patients after 24h, by enzyme-linked immunoassay (BMS213INST, eBioscience) according to manufacturer’s instructions. Supernatants were diluted 1:1000 for this protocol. The median concentration of IL-6 was 4340 and 6068 pg/ml for GCA and PMR supernatants respectively, without statistical significance between the 2 groups (p>0.99).

**Sera experiments**

Having already confirmed the induction of cellular senescence by GCA tissue culture supernatant, we tested the potential impact of sera derived from GCA patients (n=3) and PMR patients (n=3) on cells. Soluble level IL-6 was quantified and sera were applied for 24h on primary skin fibroblasts diluted (dilution 1/10). GL13 and IL-6 were not detected in both settings after treatment.

**Tissues**

Approximately 1.5-2 cm of temporal artery was received per patient and was cut transversely every 3 mm; each tissue segment had thickness of 0.2 mm. Fresh (20 patients) and Formalin-Fixed and Paraffin Embedded (FFPE) (all patients) artery tissues were used. Surgical removal of temporal arteries was performed by the same surgeon in each center following the same procedure.

**Artery tissue cultures**

The *ex vivo* short-term artery tissue culture system was developed as described previously [1]. Briefly, fresh biopsy slices of 5mm were cultured in full DMEM medium at 37^o^C and 5% CO_2_ for 24 hours, followed by the collection of cultured conditioned media containing the inflammatory micro-environment of the inflamed tissue. We have collected and stored at -80^o^C the 24h cultured conditioned supernatant of: i) temporal arteries from of 10 GCA and 10 PMR patients, ii) 1 abdominal aortic aneurism surgically excised for therapeutic purposes from 1 GCA patients with aortitis and iii) 2 normal arteries from non-vasculitis individuals serving as negative controls. Optimization of all cultures has been performed by evaluating: a) tissue cell viability and b) IL-6 levels at 24h culture conditions.

**Cell line - *Ex vivo* experiments**

Primary skin fibroblasts derived from a healthy donor (fifteen years old) after written consent, were used for the *ex vivo* experiments. Cells were maintained in 15% Fetal Bovine Serum (F7524, Sigma-Aldrich)/ 1% (v/v) pen/strep (L0022V, Biowest/ DMEM medium (L0102, Biowest) at 37°C and 5% CO_2_. 7.500 cells were seeded on coverslips per well in a 24 well plate. Cells were incubated with either tissue culture supernatants (dilution 1/2, total volume 400μl) or patients’ serum (dilution 1/10, total volume 400μl) for 5 days according to our preliminary experiments that demonstrated the occurence of senescent cells at this time point (**Supplementary Figure 3**). Interleukin-6 receptor (IL-6R) and Interleukin 1-β receptor (IL-1βR) were blocked in primary skin fibroblast before treatment with artery tissue supernatant, using 1 μg/ml of Tocilizumab (RoActemra, Roche) and 10 μg/ml of Anakinra (Kineret*, Swedish Orphan Biovutrum AB) respectively, as described previously [2, 3]. Fibroblasts were pre-treated for 2 hours with ILs-R inhibitors and then tissue culture supernatant were applied for 5 days. Presence of inhibitors was maintained through all days of the experiments. After 5 days, coverslips were washed with PBS and fixed with 4% Paraformaldehyde (10 minutes, 4^o^C). Coverslips were washed with PBS and stored at 4^o^C.

**Antibodies**

The following antibodies were used in the current study for immunohistochemistry and immunofluorescence studies. Primary antibodies: anti-biotin (1/300, ab201341, abcam), anti-biotin (1/100, BNC610400-100, Biotium), anti-p21^WAF/Cip1^ (#2947, Cell Signaling Technology), anti-IL-6 (1/200, MAB206, R&D systems), anti-aSMA (1/100, ab5694, abcam) anti-Vimentin (1/500, ab92547, abcam), anti-CD34 (1/200, ab81289, abcam), anti-CD68 (1/200, ab213363, abcam), anti-MMP-9 (1/150, sc-21733, Santa Cruz). Secondary antibodies: anti-mouse (1/500, 20014, Biotium), anti-rabbit (1/1.000, 20098, Biotium).

**Immunohistochemistry - GL13**

FFPE tissues were cut in 4μm sections and stained for detection of senescent cells using GL13 reagent as described previously [4]. Briefly, sections were deparaffinized, hydrated and antigen retrieval was heat-mediated using citric acid buffer (pH=6) for 15 minutes in steamer. Samples were cooled for 20 minutes in ice bath and GL13 was applied for 2 times, 10 minutes each at 37^o^C and then positive signal was developed using the Dako REAL EnVision Detection System, (Cat.no: K5007) according to the manufacturer’s instructions. Primary anti-biotin antibody (ab201341, abcam) was applied in dilution 1:300 at 4^o^C overnight. Specimens were counterstained with Hematoxylin. GL13 was quantified at least in 3 tissue segments per patient. The whole segments were scanned and the mean percentage of GL13 positive cells was obtained from 5-10 high power fields [5]. Inflammatory cells in GCA TABs were not included in the quantification process.

**Immunofluorescence**

In this work for the first time in the literature we were able to apply successfully immunofluorescence detection of lipofuscin (GL13) in a triple staining protocol on paraffin embedded tissues. FFPE tissues were cut in 4μm sections, deparaffinized, hydrated and antigen retrieval was heat-mediated using citric acid buffer (pH=6). Section heated into citric buffer (pH=6) for 15 minutes in steamer (for GL13, anti-IL-6, anti-MMP-9, anti-Vimentin, anti-aSMA, anti-CD68 and anti-CD34 staining) or in microwave (for anti-p21^WAF1/Cip1^ staining) for 30 minutes. Samples were cooled for 20 minutes in ice bath and washed with PBS while blocking of non-specific binding sites was performed using goat-serum (1/40, ab7481, abcam) for 2 hours. After a washing step with PBS, GL13 was applied for 2 times, 10 minutes each, 37^o^C and primary anti-biotin antibody (1/100, BNC610400-100, Biotium) was applied for 1h in RT. In the case of multi-staining, sections were initially washed with PBS and incubated with the mix of primary antibodies for the respective markers for 1h in RT; subsequently they were washed with PBS and incubated with the mix of secondary antibodies for 1h in RT. Tissue autofluorescence was diminished by applying Vector® TrueVIEW® Autofluorescence Quenching kit (SP-8400-15). Samples were washed with PBS and stained with DAPI, washed again with dH_2_0 and mounted. For immunofluorescence staining on fixed cells, coverslips were treated with 0.3% Triton/PBS (15 minutes) for cellular membrane permeabilization. Coverslips were washed and incubated with goat-serum for blocking of non-specific binding sites (1/40, ab7481, abcam) for 2 hours in RT. After cells washing, GL13 was applied for 10 minutes 37^o^C. Primary anti-biotin antibody (1/100, BNC610400-100, Biotium) was applied for 1h and then cells were washed. In the case of multiple staining, the mix of primary antibodies for the corresponding antigens, was applied for 1h in RT following a wash step with PBS. Then cells were incubated with the mix of secondary antibodies for 1h in RT. Cells were than washed, stained with DAPI, washed again with dH_2_O and mounted.

GL13 and p21^WAF1/Cip1^ quantification was performed as described in Immunohistochemistry section. Results for senescence cell type analysis were quantified separately for each cell population. For multi staining with vimentin, the media layer of each artery using 5 representative fields with at least 500 cells per sample was quantified. For multi staining with CD68 and CD34, quantification of the arterial endothelium and vasa vasorum quantification was performed on the whole tissue section in 5 representative fields with at least 100 cells per sample. For IL-6 and MMP-9 quantification analysis the whole tissue section was analyzed using 5-10 fields with at least 500 total cells per sample.

**Optical, Confocal and Electron Microscopy**

Tissues and cells were observed under ZEISS Axiolab 5 optical microscope using objectives 20x and 40x. Leica TCS-SP8 confocal microscope was used at objectives 20x, 40x and 63x. **Transmission electron microscopy (TEM)**

Small tissue fragments from artery biopsies of PMR and GCA were fixed in 2.5% glutaraldehyde in 0.01M phosphate buffer saline (PBS) for 3h at RT and postfixed in 1% aqueous osmium tetroxide (OsO4) for 1h at 4⁰C. The specimens were then dehydrated through a graded series of ethanol, infiltrated gradually in a mixture of Epon/Araldite resins diluted in propylene oxide (1:2, 1:1, 2:1, 1 h each) and finally embedded in fresh epoxy resin mixture. Subsequently, ultrathin sections (80-90nm thickness) of epoxy-embedded specimens were cut with a Diatome diamond knife on an ultramicrotome Leica Ultracut R. The sections were mounted onto 200 mesh copper grids, stained with ethanolic uranyl acetate and lead citrate and finally, were observed in a FEI Morgagni 268 transmission electron microscope. Electron micrographs were taken with an Olympus Morada digital camera.

**RNA-seq bioinformatic analysis**

Bulk RNAseq analysis in already published data from TABs of GCA and non GCA patients was conducted to investigate the possible involvement of genes and pathways associated with cellular senescence. Raw counts of 40 FFPE temporal artery (TA) sections that were collected from biopsy-positive GCA participants older than 50 years old and 31 normal TAs (without herpes zoster, diabetes, cancer, substance abuse, or immunosuppression) that were removed postmortem from participants over than 50 years of age, were downloaded from Gene Expression Omnibus with accession number GSE174694 [6] and used for further analysis. Normalization of reads and removal of unwanted variation was performed with RUVseq [7]. Differentially expressed genes were identified using the DESeq 2 [8] R package and genes with log2 fold change cut-off of 1 and p-adjust less than 0.05 were considered to be significant. Gene ontology and pathway analysis was performed using the Database for Annotation, Visualization and Integrated Discovery (DAVID) [9]. Only pathways and biological processes with p-value lower than 0.05 were considered to be significantly enriched.

1.	O'Neill L, Rooney P, Molloy D, Connolly M, McCormick J, McCarthy G, et al. Regulation of Inflammation and Angiogenesis in Giant Cell Arteritis by Acute-Phase Serum Amyloid A. Arthritis Rheumatol. 2015 Sep; 67(9):2447-2456.

2.	Al-Jomah N, Al-Mohanna FH, Aboussekhra A. Tocilizumab suppresses the pro-carcinogenic effects of breast cancer-associated fibroblasts through inhibition of the STAT3/AUF1 pathway. Carcinogenesis. 2021 Dec 31; 42(12):1439-1448.

3.	Nicolas AM, Pesic M, Engel E, Ziegler PK, Diefenhardt M, Kennel KB, et al. Inflammatory fibroblasts mediate resistance to neoadjuvant therapy in rectal cancer. Cancer Cell. 2022 Feb 14; 40(2):168-184 e113.

4.	Evangelou K, Lougiakis N, Rizou SV, Kotsinas A, Kletsas D, Munoz-Espin D, et al. Robust, universal biomarker assay to detect senescent cells in biological specimens. Aging Cell. 2017 Feb; 16(1):192-197.

5.	Evangelou K, Veroutis D, Paschalaki K, Foukas PG, Lagopati N, Dimitriou M, et al. Pulmonary infection by SARS-CoV-2 induces senescence accompanied by an inflammatory phenotype in severe COVID-19: possible implications for viral mutagenesis. Eur Respir J. 2022 Aug; 60(2).

6.	Bubak AN, Mescher T, Mariani M, Frietze SE, Hassell JE, Jr., Niemeyer CS, et al. Targeted RNA Sequencing of Formalin-Fixed, Paraffin-Embedded Temporal Arteries From Giant Cell Arteritis Cases Reveals Viral Signatures. Neurol Neuroimmunol Neuroinflamm. 2021 Nov; 8(6).

7.	Risso D, Ngai J, Speed TP, Dudoit S. Normalization of RNA-seq data using factor analysis of control genes or samples. Nat Biotechnol. 2014 Sep; 32(9):896-902.

8.	Love MI, Huber W, Anders S. Moderated estimation of fold change and dispersion for RNA-seq data with DESeq2. Genome Biol. 2014; 15(12):550.

9.	Huang da W, Sherman BT, Lempicki RA. Systematic and integrative analysis of large gene lists using DAVID bioinformatics resources. Nat Protoc. 2009; 4(1):44-57.

**References**
